# A High HIV-1 Strain Variability in London, UK, Revealed by Full-Genome Analysis: Results from the ICONIC Project

**DOI:** 10.1101/139642

**Authors:** Gonzalo Yebra, Dan Frampton, Tiziano Gallo Cassarino, Jade Raffle, Jonathan Hubb, R Bridget Ferns, Zisis Kozlakidis, Andrew Hayward, Paul Kellam, Deenan Pillay, Duncan Clark, Eleni Nastouli, Andrew J. Leigh Brown, on behalf of the ICONIC consortium

**Affiliations:** Institute of Evolutionary Biology, University of Edinburgh, Edinburgh, UK; UCL Division of Infection and Immunity, Faculty of Medical Sciences, London, UK; Department of Clinical Virology, UCL Hospital NHS Foundation Trust, London, UK; Department of Virology, Barts Health NHS Trust, London, UK; NIHR UCLH/UCL Biomedical Research Centre, London, UK; UCL Institute of Disease Informatics, Farr Institute of Health Informatics Research, London, UK; UCL Institute of Epidemiology and Health Care; Division of Infectious Diseases, Department of Medicine, Imperial College London, UK; Department of Population, Policy and Practice, UCL GOS Institute of Child Health, London, UK

## Abstract

**Background & Methods:** The ICONIC project has developed an automated high-throughput pipeline to generate HIV nearly full-length genomes (NFLG, i.e. from *gag* to *nef*) from next-generation sequencing (NGS) data. The pipeline was applied to 420 HIV samples collected at University College London Hospital and Barts Health NHS Trust (London) and sequenced using an Illumina MiSeq at the Wellcome Trust Sanger Institute (Cambridge). Consensus genomes were generated and subtyped using COMET, and unique recombinants were studied with jpHMM and SimPlot. Maximum-likelihood phylogenetic trees were constructed using RAxML to identify transmission networks using the Cluster Picker.

**Results:** The pipeline generated sequences of at least 1Kb of length (median=7.4Kb) for 375 out of the 420 samples (89%), with 174 (46.4%) being NFLG. A total of 365 sequences (169 of them NFLG) corresponded to unique subjects and were included in the down-stream analyses. The most frequent HIV subtypes were B (n=149, 40.8%) and C (n=77, 21.1%) and the circulating recombinant form CRF02_AG (n=32, 8.8%). We found 14 different CRFs (n=66, 18.1%) and multiple URFs (n=32, 8.8%) that involved recombination between 12 different subtypes/CRFs. The most frequent URFs were B/CRF01_AE (4 cases) and A1/D, B/C, and B/CRF02_AG (3 cases each). Most URFs (19/26, 73%) lacked breakpoints in the PR+RT *pol* region, rendering them undetectable if only that was sequenced. Twelve (37.5%) of the URFs could have emerged within the UK, whereas the rest were probably imported from sub-Saharan Africa, South East Asia and South America. For 2 URFs we found highly similar *pol* sequences circulating in the UK. We detected 31 phylogenetic clusters using the full dataset: 25 pairs (mostly subtypes B and C), 4 triplets and 2 quadruplets. Some of these were not consistent across different genes due to inter- and intra-subtype recombination. Clusters involved 70 sequences, 19.2% of the dataset.

**Conclusions:** The initial analysis of genome sequences detected substantial hidden variability in the London HIV epidemic. Analysing full genome sequences, as opposed to only PR+RT, identified previously undetected recombinants. It provided a more reliable description of CRFs (that would be otherwise misclassified) and transmission clusters.

## Introduction

Human immunodeficiency virus type 1 (HIV-1) shows a great genetic diversity [1], which causes the emergence of multiple viral strains. HIV-1 is classified in four groups: M, O, N and P. HIV-1 group M drives the global pandemic, and includes 9 subtypes (A-D, F-H, J, K), at least 88 circulating recombinant forms (CRFs) [2] and countless unique recombinant forms (URFs), generated when cells are superinfected by HIV-1 virions from different subtypes/CRFs. CRFs are defined as inter-subtype recombinant lineages that derived from the same common ancestor, and for which at least three epidemiologically unlinked variants are monophyletic and share an identical genetic structure along their full genome. URFs are recombinants with different recombination breakpoints than those found in CRFs, and emerge after localised, isolated recombination events. URFs are responsible for at least 20% of HIV-1 infections worldwide –a proportion that reaches 40% in some African countries [3].

HIV-1 subtype B is the prevalent variant in high-income areas like North America and West Europe [3], and the most studied strain despite only accounting for 12% of global infections. But in the last decade, the remaining subtypes and recombinants (“non-B variants”, responsible for the remaining 88% of infections worldwide) are increasing in prevalence and heterogeneity in developed countries [4]. The UK is no exception, and viral genetic diversity (especially involving novel recombinants) markedly increased in the British HIV-1 epidemic between 2002 and 2010 [5]. This new heterogeneity has resulted in the description of an increasing number of CRFs in Western Europe, which traditionally were restricted to regional hotspots with great co-circulation of variants –like sub-Saharan Africa, South-East Asia and certain parts of South America. These new European recombinants include CRF47_BF and CRF73_BG in Spain and Portugal [6, 7], CRF50_A1D in the UK [8], CRF56_cpx in France [9], and CRF60_BC in Italy [10], as well as second-generation complex recombinant strains [11].

The genetic variability derived from the presence of non-B variants could affect management of HIV-1 infection [12], including the antiretroviral response [13], the efficacy of viral load and resistance assays [14], the speed of disease progression [15, 16], the emergence of drug resistance mutations [17] and the vaccine development [18]. Additionally, HIV-1 recombination is a powerful evolutionary force that occurs ten times more frequently than base substitutions [19] and produces and maintains further variability by shuffling and combining viral genetic material. HIV-1 recombination can accelerate the adaptation to the host [20], improve viral replication [21] and provide mechanisms to escape both the immune response and antiviral treatments [22-24]. At the same time, gene reshuffling derived by recombination can compensate the loss in viral fitness derived from these adaptations [20] HIV will continue to evolve and diversify, which will bring new challenges to prevention and treatment efforts.

Thus, the proper detection and description of HIV-1 variants in representative cohorts is essential, especially considering the worldwide emergence and transmission of recombinants. The coincidence of the increasing HIV-1 complexity in Western countries with the rapid development of next-generation sequencing (NGS) technology [25] highlights the potential of applying these techniques to routine surveillance of HIV-1 molecular epidemiology. Here, we aimed to identify URFs in a dataset consisting on HIV-1 nearly full-genome sequences obtained through NGS of samples taken in two major hospitals in London, UK. We describe their recombination patterns using a combination of bioinformatics tools, which include bootscanning and phylogenetic analysis of sequence sub-segments, considered the reference methodology for detecting HIV-1 variants since its first use [26, 27].

## Methods

### Pipeline and study subjects

Within the ICONIC project, we have developed an automated and customisable high-throughput analysis pipeline to generate HIV clinical whole genomes, quantify minority variants and identify drug-resistance mutations (DRMs). Via de novo assembly of contigs, concatenation and correction by alignment to reference and subsequent remapping of reads, this pipeline derives a consensus sequence and minority variants across each genome [28].

In order to generate HIV consensus genomes, the pipeline was applied to 420 HIV samples, collected between 2012 and 2014 at two major London hospitals: University College London Hospital (n=332) and Barts Health NHS Trust (n=88). They were sequenced using an Illumina MiSeq at the Wellcome Trust Sanger Institute (Cambridge).

### Subtyping and recombination analyses

The consensus genomes were preliminary subtyped using a local implementation of COMET [29]. This method was applied to the full dataset but also independently to the principal structural genes (gag, *pol* and *env*), extracted using GeneCutter (*https://www.hiv.lanl.gov/content/sequence/GENE_CUTTER/* cutter.html). This approach was followed in order to detect putative recombinant, i.e. those cases in which different genes disagreed in subtype or any of them yielded an unassigned result. Those cases were analyse in more depth using the jpHMM [30] online tool, which is capable of detecting and locating recombination breakpoints in HIV-1 sequences.

In those sequences where these tools confirmed the presence of recombination events, further analyses were applied using the program SimPlot version 3.5.1 [31], which applies a bootscanning method. This consists of a sliding-window phylogenetic bootstrap analysis of the query sequence (using a window size of 300bp moving in 10bp increments) aligned against a set of reference strains to reveal breakpoints, which was later confirmed by performing phylogenetic analysis using RAxML (GTR+Γ substitution model with 1000 bootstrap replicates) of the inter-breakpoint segments. For these, we incorporated sequences from the 2010 HIV subtype reference alignment downloaded from Los Alamos National Laboratory HIV database website [1].

In addition, to study the most likely geographical origin of the parental variants that generated the URF detected, we looked for genetically similar sequences from GenBank (retrieved using HIV BLAST (*https://www.hiv.lanl.gov/content/sequence/BASIC_BLAST/basic_blast.html*)*) for each recombinant segment.*

Finally, we extracted the ‘clinical’ PR+RT *pol* segment (covering the protease and the first 235 codons of the reverse transcriptase) of the URF sequences that presented breakpoints in this coding region to retrospectively find highly genetically similar sequences within the UK HIV Drug Resistance database (UKHDRD) through BLAST –these sequences could represent recombinant lineages already circulating in the UK.

### Analysis of intra-subtype recombination

Whereas inter-subtype recombination can be detected relatively easily, intra-subtype recombination is much more difficult to identify due to a greater sequence similarity between sequences of the same subtype/variant, and the lack of standardised methodology. For the largest pure subtype datasets, we aimed to detect intra-subtype recombination using the program RDP version 4 [32] following the approach described by Kiwelu and colleagues [33]. In short, RDP4 uses 7 non-parametric recombination detection methods: RDP, GENECONV, modified Bootscan, MAXCHI, CHIMAERA, SISCAN and 3SEQ. For each of these methods, the default RDP4 settings were used. The recombination events considered definitive were those with statistically significant support (p<0.05 after Bonferroni correction for multiple comparisons) for at least 3 out of the 5 algorithms. RDP4 screens all possible sequence triplets/quartets to find recombinants and potential parents, which makes possible the analysis without reference sequences.

### Analysis of phylogenetic clusters

In order to identify potential transmission clusters in the full dataset, we constructed maximum likelihood trees with RAxML (with the GTR+Γ substitution model) and applying the Cluster Picker with a cutoff of 90% for node statistical support and ≤6% for the intra-cluster genetic diversity. In order to minimise the effect of recombination in the detection of clusters, further Cluster Picker analyses of the genes gag (1.5Kb), *pol* (3Kb) and *env* (2.5Kb) were performed, using genetic diversity cutoffs of ≤5.5%, ≤4.5% and ≤9%, respectively. These levels were chosen taking as reference the value commonly used for *pol* [34] and adjusting the rest of the genes according to their different evolutionary rates.

### Ethics statement

The ICONIC project (IRAS project ID 131373) was approved by the NHS Research Ethics Committee (13/LO/1303). ICONIC was also included in the Portfolio Database of the UK Clinical Research Network (UKCRN ID: 15776). The ICONIC study does not require written informed consent by study participants as it only uses pseudonymised residual diagnostic material and no patient genetic material.

## Results

### Sequence dataset

The pipeline generated sequences of at least 1Kb of length (median=7.4Kb) for 375 out of the 420 ICONIC samples (89%), with 174 of the 375 (46.4%) sequences covering the full ‘clinical’ genome (i.e. from *gag* to *nef*). The rest of sequences (n=201) had variable coverage. Of the 375 sequences to be analysed, 365 corresponded to unique patients and were included in the subsequent analyses. The 375 assembled genomes were deposited in GenBank under the accession numbers MF109352 to MF109726 (see **Supplementary Table 1**). All the raw paired-end short reads were deposited in the European Nucleotide Archive (ENA) under the study accession number PRJEB6008.

### HIV-1 subtype distribution

The most frequent HIV strains were subtypes B (n=149, 40.8%) and C (n=77, 21.1%) and CRFs 02_AG (n=32, 8.8%) and 01_AE (n=15, 4.1%). In total, we found 14 different CRFs (n=66, 18.1%). We also found multiple URFs (n=32, 8.8%) that involved recombination between 12 different parental variants: 7 subtypes (A1, B, C, D, F1, G and J) and 5 CRFs (CRF01_AE, CRF02_AG, CRF06_cpx, CRF22_01A1 and CRF50_AD). The most frequent URFs were B/CRF01_AE (4 cases) and A1/D, B/C, and B/CRF02_AG (3 cases each). Their recombination patterns are shown in **Supplementary Figure 1**.

**Figure 1.**
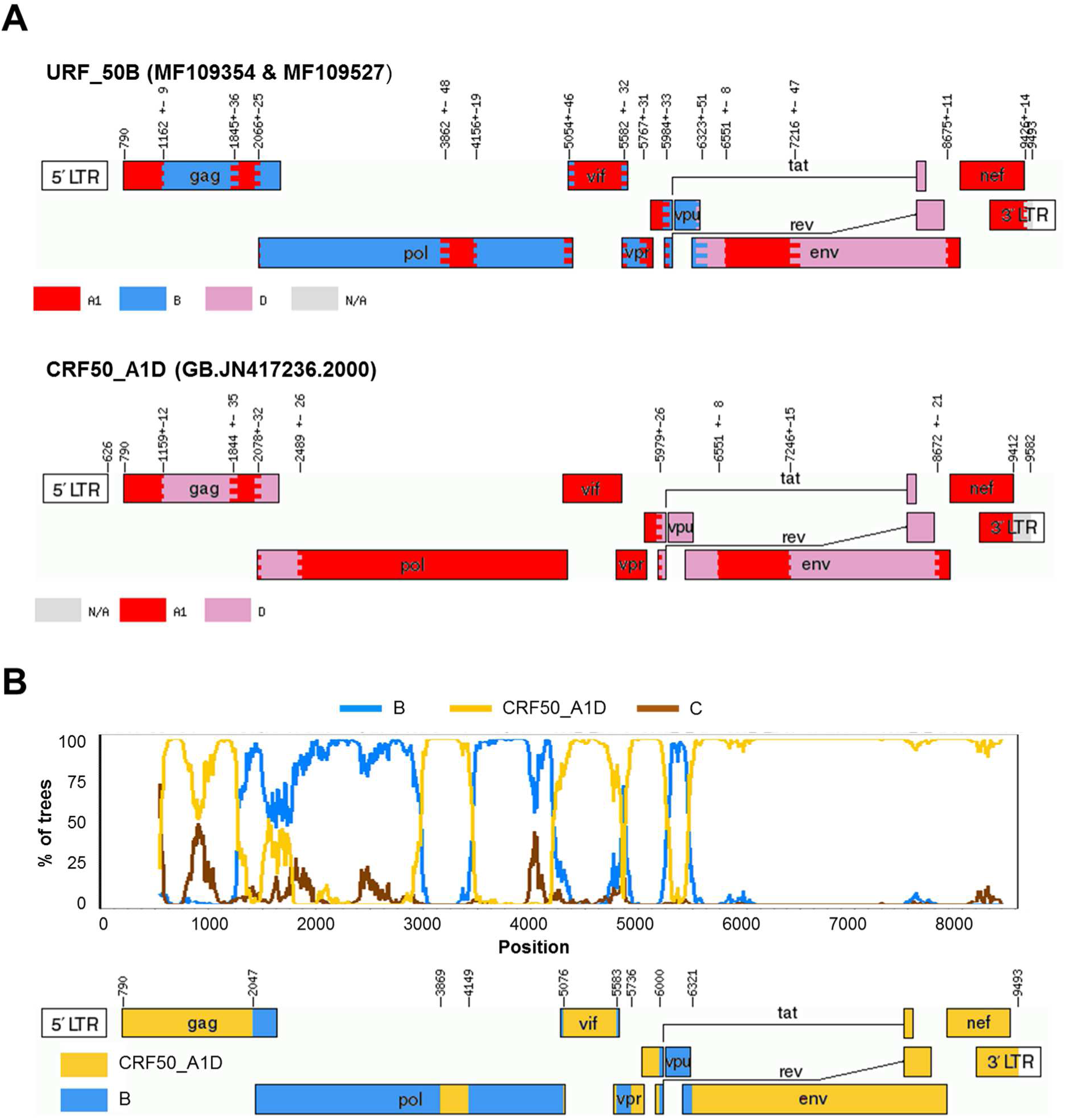
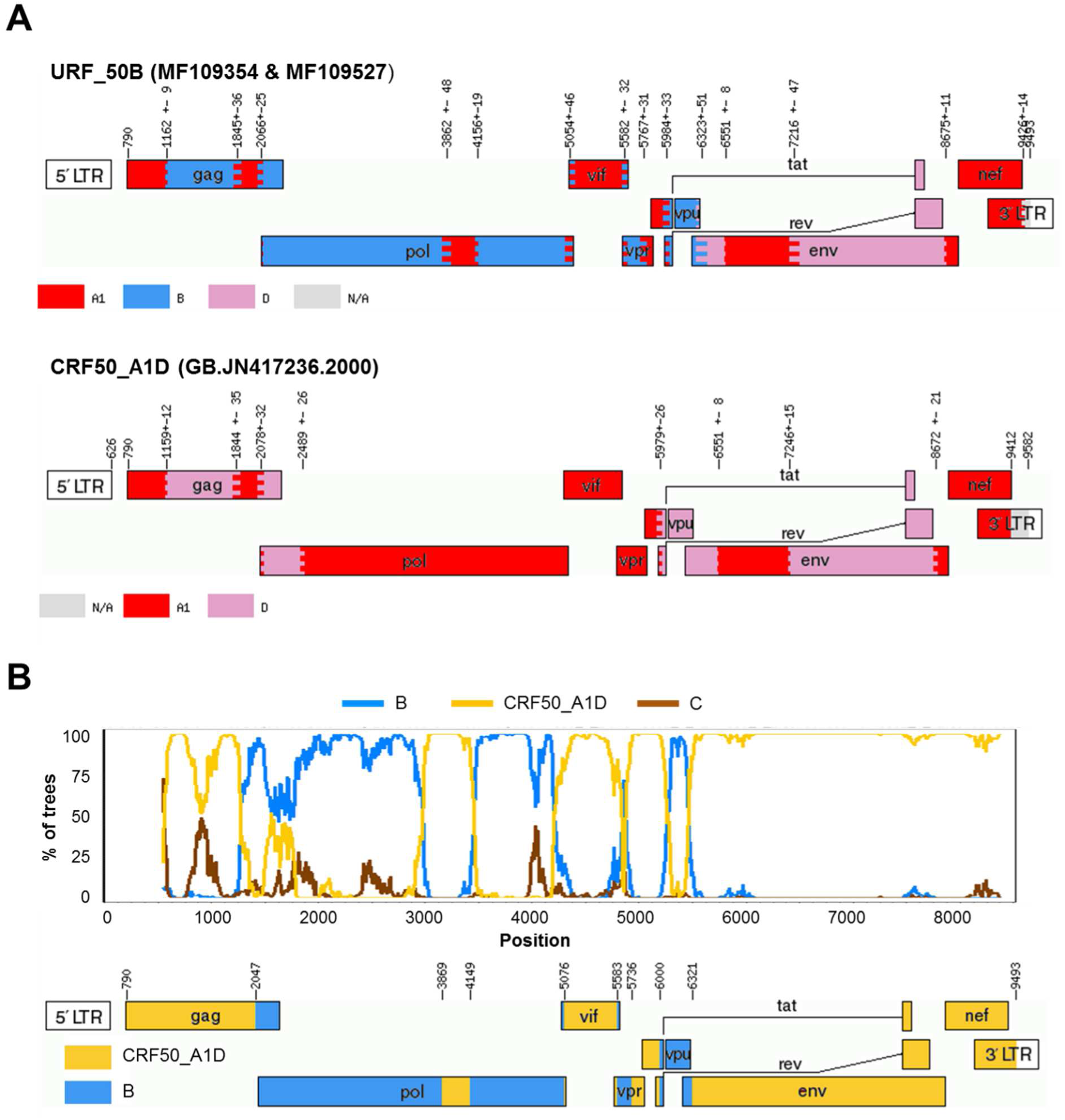
A) Recombination pattern (according to jpHMM) of the two URF_50B (accession numbers MF109354 and MF109527) discovered in the ICONIC dataset and the GenBank full-length sequence JN417236, prototype of the recombinant CRF50_A1D. B) Bootscanning analysis of the URF_50B performed using SimPlot and diagram showing the definitive recombination pattern.

Importantly, we generated new sequences of several CRFs that remain poorly (n<10) represented in public sequence databases, providing full-genome each of the following recombinants: CRF03_AB (only 3 genome sequences available in GenBank), CRF13_cpx (8), CRF18_cpx (5), CRF25_cpx (6) and CRF43_02G (4). Also, we provide partial genomes of CRF09_cpx, CRF19_cpx, CRF49_cpx and CRF60_BC (one each), which will provide new and invaluable material for the study of these recombinant lineages.

### Subtype distribution using partial *pol*

In order to explore the implications of using only the PR+RT partial *pol* sequence to study the HIV-1 subtype distribution, we extracted the 277 sequences (out of the total 365 sequence dataset, 75.9%) that covered this region and applied the COMET and SCUEAL subtyping tools. SCUEAL was used to make the results comparable to those from the UKHDRD reports [5]. The prevalence of pure subtypes in the *pol* gene (74.4% for COMET and 73.0% for SCUEAL) was similar to that found in the full-length ICONIC dataset (73.2%), with the only exception of an overestimation of subtype G by SCUEAL (7.0% using *pol* versus 2.7% in the definitive results; chi-square test *P*=0.01). Differences found in the proportion of recombinants (CRFs and/or URFs) were also not significant.

Focusing on the 32 URF sequences, 26 covered the 1.3Kb PR+RT *pol* region, of which only 7 (7/26, 26.9%) presented a breakpoint in that region and were consequently identified as recombinants by both COMET and SCUEAL. The remaining 19 (19/26, 73%) URFs were therefore mostly considered as pure subtypes/CRFs when only the PR+RT *pol* sequence was analysed. As the PR+RT fragment represents only ∼15% of the size of the genomes analysed here, it is in fact detecting more recombinants than expected (26.9% versus 8.8% when analysing the genome data; chi-square test *P*=0.008).

### Unique recombinant forms (URFs)

The 32 URF sequences were further explored by analysing the parental lineages that generated them and their closest sequences in public sequence databases identified by BLAST. For those URF presenting breakpoints in the PR+RT *pol* region, genetically similar *pol* sequences were searched within the UK HIV Drug Resistance database (UKHDRD). In the following paragraphs we provide a detailed explanation for the most frequent and/or complex URF. The rest of the URF analyses are detailed in the **Supplementary Results.**

**B/CRF50_A1D** pair (accession numbers MF109354and MF109527, respectively). These two URF_50B sequences that formed a phylogenetic pair (genome-wide similarity=96.2%) were classified as an A1/B/D recombinant by jpHMM, but after phylogenetic analysis of the recombinant segments, those assigned to A1 and D clustered with the recombinant CRF50_A1D (previously described in the UK [8]), and shared the same breakpoints between A1 and D found in this CRF (**Figure 1**). Therefore, they were reclassified as B/CRF50_A1D. The subtype B segment that covered the PR+RT region in *pol* presented high similarity (>98%) to UK “pure” subtype B sequences in both the UKHDRD dataset and GenBank, which suggests that these recombinants originated in the UK. A different URF_50B, presenting different breakpoints as those reported here, was already described in Foster et al. [8] (see below for more information on this recombinant sequence), suggesting that this CRF of Eastern African origin is actively recombining with local sequences.

**A1/B**. Two URF_A1B were identified, with one of them (MF109544) presenting a long gap between the A1 and B portions. The A1 segment was closest to sequences from Kenya (similarity=93%), whereas no close GenBank sequences to the subtype B portion could be found. The other URF_A1B found (MF109371) presented a small A1 fragment in *pol* (694bp-long) in a background of subtype B. Whereas the latter did not present high similarity close to any GenBank sequence in *gag* or *env*, the A1 fragment was intimately related to CRF50_A1D in the phylogenetic analysis, and particularly (similarity=95%) to a B/CRF50_A1D recombinant sequence (accession number JN417238, see above) also described in the article by Foster and colleagues [8] where this CRF is described. JN417238 and this newly described URF_A1B shared common A1/B breakpoints in the PR+RT region although the latter presented additional breakpoints absent in the URF_A1B; however, they did not cluster together in their common subtype B fragments. The PR+RT *pol* portion of both the URF_A1B described here and, especially, of JN417238 were genetically similar (98%) to an A1/B cluster already described in Canada that was identified through BLAST; we also found 5 GenBank A1/B sequences sampled in the UK which were very similar. Within the UKHDRD, we identified 13 more PR+RT *pol* sequences circulating sharing breakpoints with this URF_A1B according to jpHMM. These 13 UKHDRD sequences were mostly sampled on 2011-2012 but the oldest one was sampled in 2006; and in all cases with available information, the patients had been born in the UK. According to the ML tree (**Figure 2**), this A1/B lineage (and particularly JN417238, the URF_50B previously described) initiated the Canadian cluster that affected men-having-sex-with-men (MSM) population [35]. The clade also included 2 GenBank sequences from Albania and Korea, respectively, which lied basal to the UK/Canadian cluster, and points to a further international spread of this clade.

**Figure 2 (previous page).**
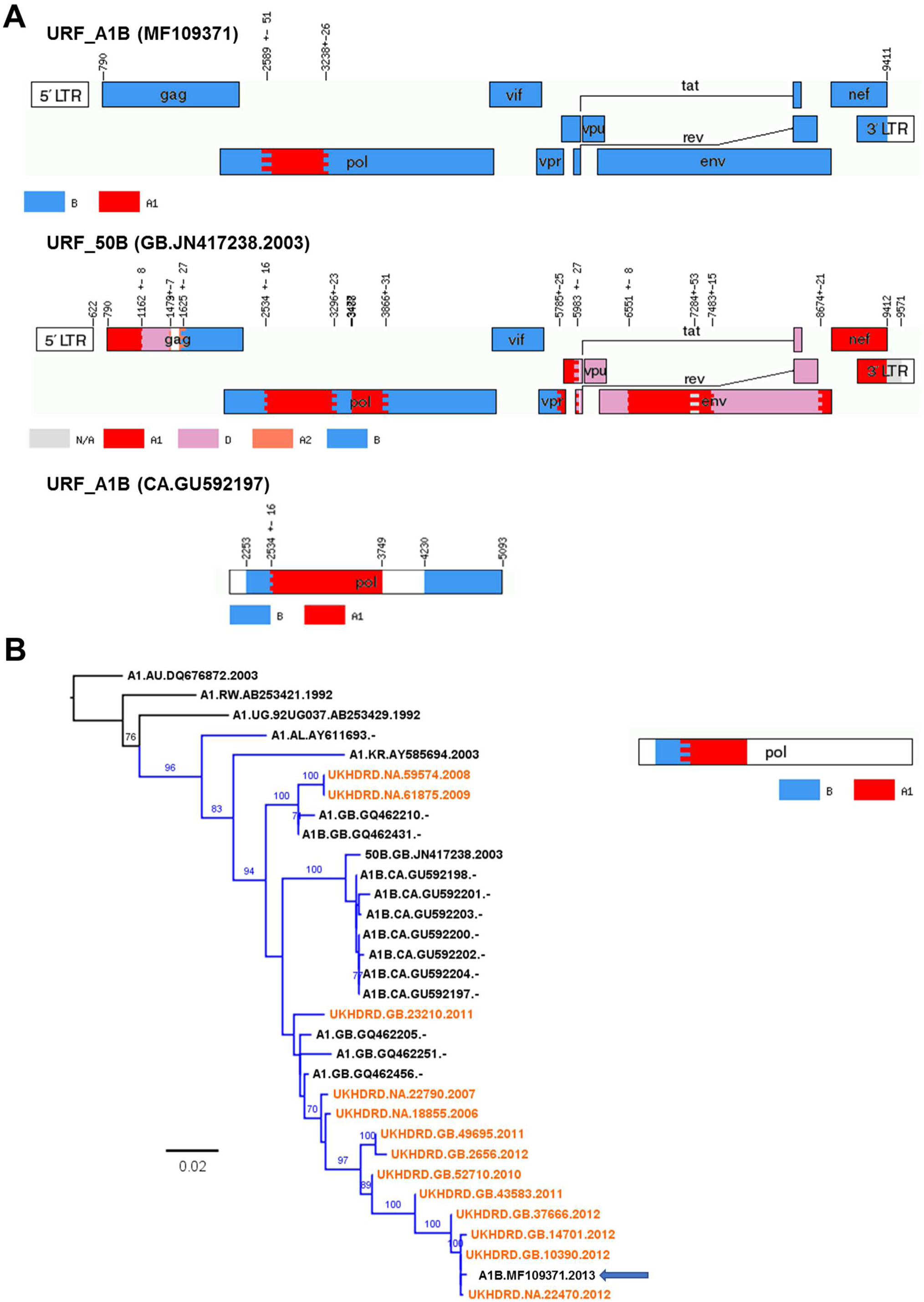
Panel A) Recombination pattern (according to jpHMM) of the URF_A1B (accession number MF109371) discovered in the ICONIC dataset, the URF_50B full-length sequence (accession number KT022394) described by Foster and colleagues and a member of the A1/D Canadian cluster described by Brenner and colleagues (accession number GU592197). These 3 which presented high similarity in the PR+RT *pol* region. B) Maximum-likelihood tree of the PR+RT *pol* region (1,000bp) of the A1/B cluster of sequences with high similarity to the ICONIC URF_A1B. Tips corresponding to the UKHRD are in orange. Tip names show, for the GenBank sequences: subtype, sampling country, accession number and sampling year (not available for the Canadian sequences); for UKHDRD sequences: country of birth, ID and sampling year. Branches corresponding to the A1/B cluster are shown in blue. The ICONIC URF_A1B sequence is highlighted with an arrow. The tree is rooted to subtype A1 reference sequences. The diagram at the right shows the recombination pattern shown by the sequences in the cluster, according to jpHMM.

**A1/D**. We detected three URF_A1D sequences (MF109466, MF109548 and MF109564) with different recombination patterns. In all cases, the closest sequences in GenBank were sampled in Eastern Africa (Uganda, Kenya and/or Tanzania). Only one of them (MF109466) presented a recombination breakpoint in the PR+RT *pol* region, for which through BLAST we identified 27 *pol* sequences in the UKHDRD and 21 in GenBank (from Kenya, Uganda and Tanzania, up to 96% of similarity) which showed a very similar A1/D breakpoint in the PR+RT region. The 27 UKHDRD sequences corresponded exclusively to patients born in Uganda, Kenya and Zimbabwe, and their samples were taken from 2001 to 2011. Both sets of sequences sampled in the UK and GenBank sequences sampled in Eastern African countries were interspersed in the tree (**Figure 3**) suggesting multiple introductions to the UK. One of the GenBank sequences from Kericho, Kenya (accession number KT022394) had a full-length sequence available which showed a recombination pattern that was very similar to that of this URF_A1D across the genome in the overlapping regions, only different at the 3’ end of the *env* gene. The presence of two sequence gaps in our sample prevented further exploration of the relatedness of these 2 sequences.

**Figure 3 (previous page).**
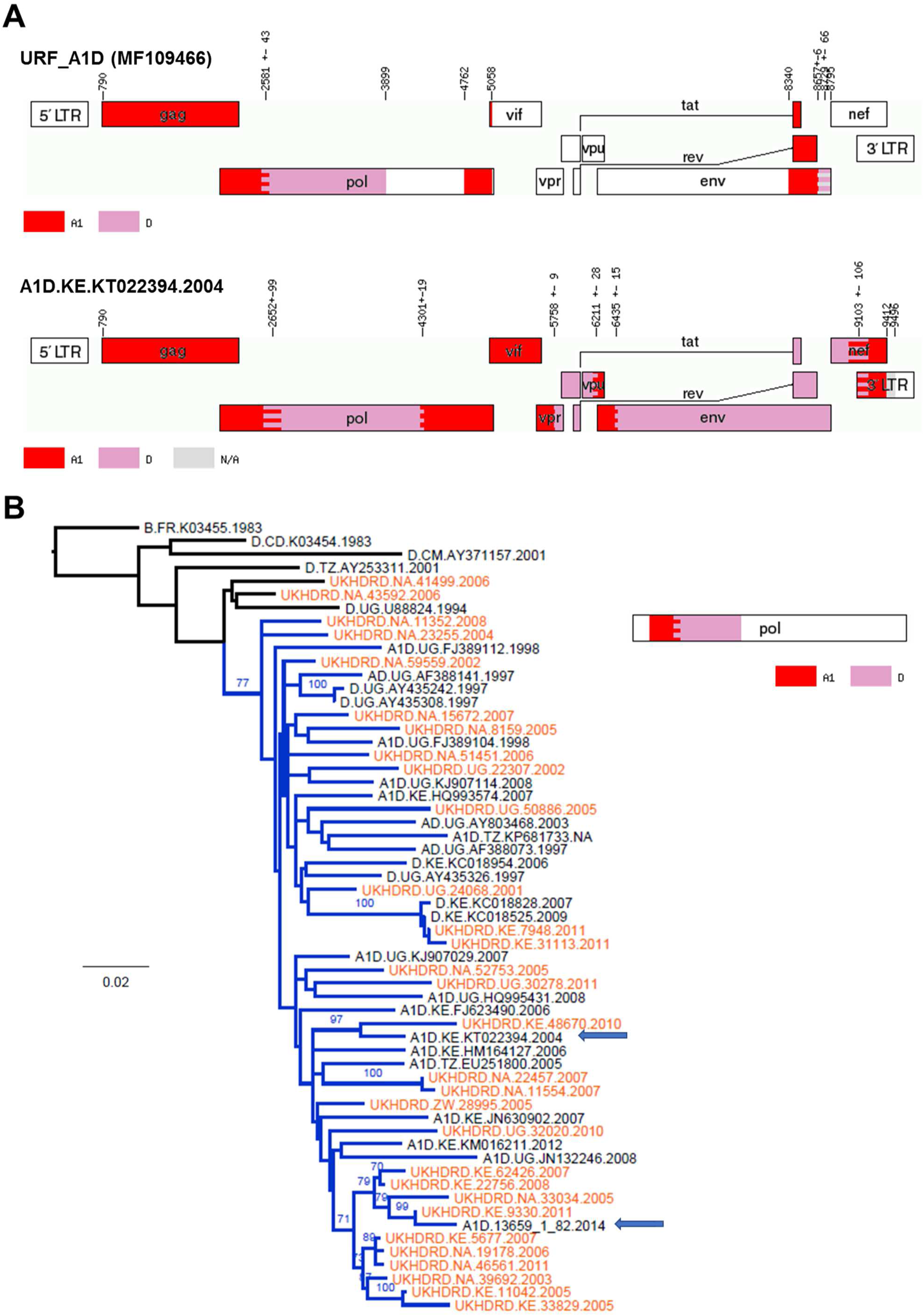
Panel A) Recombination pattern (according to jpHMM) of the URF_A1D (accession number MF109466) discovered in the ICONIC dataset and the GenBank full-length A1/D sequence KT022394, which presented high similarity in the PR+RT *pol* region. B) Maximum-likelihood tree of the PR+RT *pol* region (1,000bp) of the A1/D cluster of sequences with high similarity to the ICONIC URF_A1D. Tips corresponding to the UKHRD are in orange. Tip names show, for the GenBank sequences: subtype, sampling country, accession number and sampling year; for UKHDRD sequences: country of birth, ID and sampling year. Branches corresponding to the A1/D cluster are shown in blue. The ICONIC URF_A1D and the GenBank sequence KT022394 are highlighted with arrows. The tree is rooted to a subtype B sequence. The diagram at the right shows the recombination pattern shown by the sequences in the cluster, according to jpHMM.

**CRF06_cpx/CRF22_01A1**. We found 1 URF_0622 (MF109682) which was initially identified as a recombinant between subtypes A1, G and J and CRF01_AE by jpHMM. However, further analysis with SimPlot, which was confirmed by phylogenetic analysis of the recombinant segments, revealed that segments ascribed to subtype J and G that partially covered from *vif* to *env* actually corresponded to CRF06_cpx, and that the segments classified as A1 and CRF01_AE in *gag*, *vif* and *nef* clustered very closely to CRF22_A101 –a recombinant lineage detected in Cameroon [36] (**Figure 4**). Unfortunately, this URF_0622 sequence lacked the PR+RT *pol* region, which prevented from identifying more cases of this recombinant in the UKHDRD.

**Figure 4.**
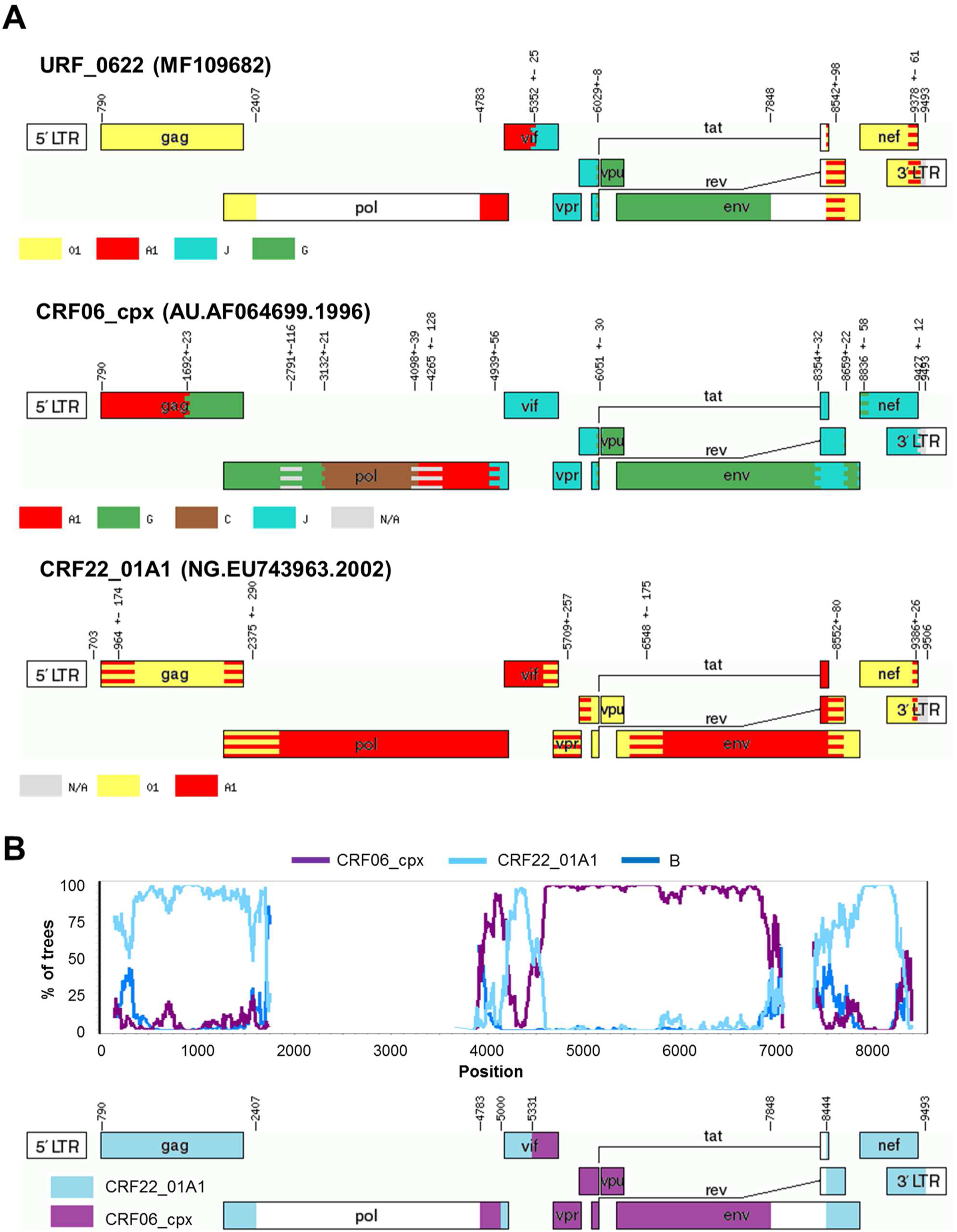
A) Recombination pattern (according to jpHMM) of the URF_0206 (15228_1_49) discovered in the ICONIC dataset and the sequences L39106 and AF064699, prototypes of the recombinants CRF02_AG and CRF06_cpx, respectively. B) Bootscanning analysis of the URF_0206 performed using SimPlot and diagram showing the definitive recombination pattern.

**Figure 4.**
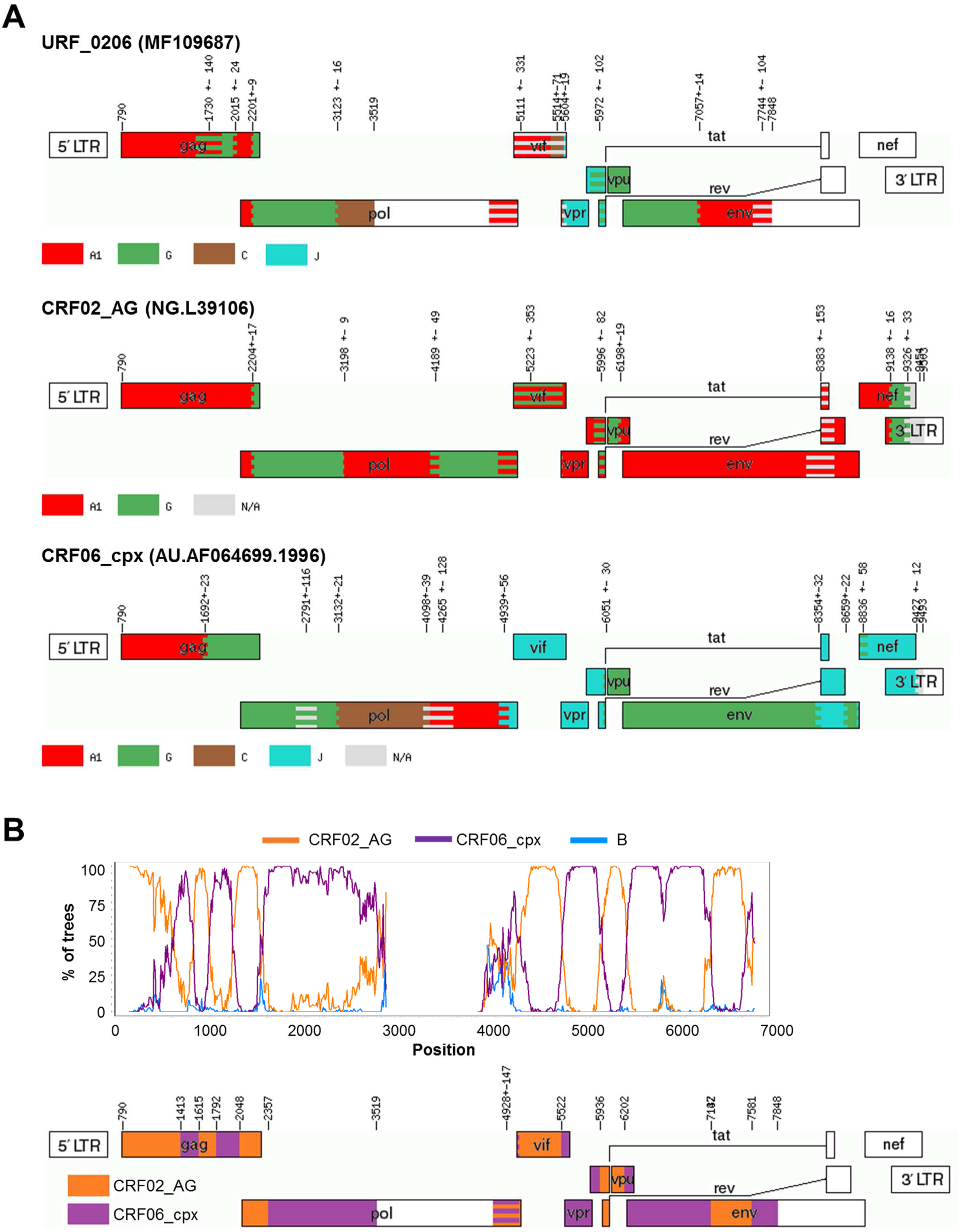
A) Recombination pattern (according to jpHMM) of the URF_0622 (accession number MF109682) discovered in the ICONIC dataset and the sequences AF064699 and EU743963, prototypes of the recombinants CRF06_cpx and CRF22_01A1, respectively. B) Bootscanning analysis of the URF_0106G performed using SimPlot and diagram showing the definitive recombination pattern.

**CRF02_AG/CRF06_cpx** (MF109687). This URF presented segments of subtypes A1, G, C and J that corresponded in ML trees and SimPlot to the recombinant CRF06_cpx (which is made up by subtypes A1, C, G and J). Interestingly, subtype A1 fragments in this URF clustered within CRF02_AG and not CRF06_cpx (**Figure 5**). In fragments in which both CRFs correspond to subtype G (in this case, the beginning of *pol* and *env* genes) the presence of further breakpoints cannot be ruled out. Recombinants between CRF02_AG and CRF06_cpx have already been discovered in Western Europe [37] probably introduced from Western Africa, a region where both CRF frequently co-circulate. Indeed, the closest GenBank sequences to this URF’s PR+RT *pol* region were “pure” CRF06_cpx sequences from Mali, Burkina Faso and Senegal (96%).

### Intra-subtype recombination

We analysed the largest ‘pure’ subtype datasets (i.e. subtypes B and C with 149 and 77 sequences, respectively) for evidence of intra-subtype recombination using RDP4. A sequence was considered an intra-subtype recombinant if the recombination signal was statistically significantly supported by at least 3 of the methods implemented in RDP4. We found 20 subtype B sequences (13.4%) that were identified as recombinants with enough statistical support. For subtype C, 7 sequences (9.1%) were intra-subtype recombinants. This difference was not statistically significant.

### Analysis of phylogenetic clusters

We detected 31 phylogenetic clusters using the full sequence dataset. Of them, 25 were pairs (corresponding mostly to subtypes B (n=11) and C (n=4)), 4 were triplets (1 each for A1, B, F1, and CRF06_cpx) and 2 quadruplets (CRF02_AG and CRF06_cpx). These clusters involved 70 sequences, 23.5% of the dataset. In order to explore the consistence of these clusters across different genes, we also examined clusters analysing the main HIV-1 genes independently: *gag*, *pol* and *env*. Seventeen (54.8%) of the 31 clusters detected in the full dataset were also found in the individual analyses of *gag*, *pol* and *env*. Their intra-subtype maximum genetic distance and bootstrap support values for each gene is shown in the **Supplementary Table 2**. The average genetic distance across the 17 clusters was 2.06% for the genome-wide analysis (range: 0.08-5.27%), 1.75% (0-5.19%) for *gag*, 1.33% (0.03-2.96%) for *pol* and 3.05% (0-8.92%) for *env*.

Most of the inter-gene cluster disagreements (6/14, 43%) were due to the fact that not all sequences in a given cluster completely covered the 3 genes. In 4 (29%) cases the sequences were too divergent (above the genetic distance cutoff) in one of the genes: *gag* and *pol* in 2 cases (genetic distances of 6.0-7.6%), *gag* alone (7.9%) in one case, *env* alone (12.9%) in the last one. The last disagreement (7%) was due to the fact that the statistical support of the node was below the cutoff (boostrap=88%) in the *gag* gene tree. Finally, there were 3 cases (21%) in which certain sequences showed entirely different tree topologies according to the gene analysed. One of them involved a phylogenetic pair of pure F1 sequences that clustered very closely with a URF_02BF1 (MF109637, whose recombination pattern is described in **Supplementary Results** and shown in **Supplementary Figure 1**) only in the *gag* gene, which suggests that the former could represent the parental lineage involved in the generation of that URF. The other 2 cases of phylogenetic inconsistence corresponded to subtype B clusters in which presumably, as the RDP analysis suggested, intra-subtype recombination had modified their clustering pattern.

## Conclusions

The analysis of HIV-1 nearly full-genome sequences generated by the ICONIC project detected substantial variability in the London HIV-1 epidemic. The HIV-1 subtype distribution in our dataset generally agrees with that reported in the last update (corresponding to 2010) for the whole UK based on analysis of *pol* sequences [5]. The main difference lies on the prevalence of CRFs, which was 6.2% in that report in contrast to the 3-fold higher 18.1% presented here. We demonstrate here how using the *pol* sequence alone, in this case using SCUEAL as Dolling and colleagues did, the proportion of CRFs was underestimated –other subtyping tools, such as COMET, will face other limitations. This highlights how the analysis of nearly full-genome sequences is key when identifying already established recombinant lineages. In addition, CRF lineages that are traditionally very infrequent in Western countries but were identified in the present study (e.g. CRF19_cpx, CRF49_cpx, CRF43_02G and CRF60_BC) are generally absent in the reference sequence datasets that are used by automatized HIV-1 subtyping tools.

On the other hand, the use of longer sequences allowed for a more refined description of novel recombinants (URFs), since the accuracy on their detection using *pol* sequences relies on the presence of breakpoints within this short (∼1Kb) genomic region. The 32 URFs that were detected using the full dataset involved recombination between 12 different parental strains, and some of them presented a very complex recombination pattern. Interestingly, 18 (56%) of the URF contained segments corresponding subtype B, the prevalent variant in Western countries. Four of these URFs (URF_01B) could have been introduced from South East Asia, where many similar recombinants have been described, and another 2 (a URF_BF1 transmission pair) from South America, where B/F1 recombinants represent a substantial proportion of the HIV-1 infections [3]. However, the remaining 12 URFs involving subtype B probably emerged in the UK (or at least in Western Europe) after introduction of non-subtype B variants which recombined with the local subtype B. The rest of the URFs (n=14, 44%) were recombinant between HIV-1 variants that co-circulate in sub-Saharan Africa, so they could have been generated before been introduced to the UK. Some of them were particularly complex, and implied further recombination between recombinant lineages (the so-called “second-generation” recombinants), a phenomenon already detected among African migrants in Western Europe [11, 37].

Matching the ICONIC sequences to the PR+RT *pol* sequences stored in the UK Drug Resistance Database allowed us to uncover new lineages of unique recombinants already circulating in the UK. Specifically, we detected an A1/B recombinant lineage circulating among UK-born patients whose oldest sequence was sampled in 2006. Another example was a larger A1/D clade that mostly involved patients born in Uganda and Kenya and sampled as far back as 2001, but also included many sequences from GenBank sampled in those Eastern African countries –which might indicate several introductions in the UK as opposed to local spread from a single one.

The analysis of phylogenetic clusters did not identify large HIV-1 transmission networks, but the largest ones discovered here corresponded to recombinant variants (2 quadruplets of CRF02_AG and CRF06_cpx respectively). We found disagreements between clusters found using different viral genes (i.e. *gag*, *pol* and *env*) that in some cases highlight the fact that recombination can obscure the genetic relationships between sequences belonging to the same transmission network. This also evidences the lack of standardisation in the cluster definition when analysing genes other than *pol*, and that will be needed with the increasing generation of full-genome datasets. In an attempt to help developing genetic distance cutoffs, we provide in **Supplementary Table 1** the values found for the clusters that were consistent across genes.

Recombination between different subtypes/CRFs happens frequently and its detection is relatively easy. However, the detection of recombinants between different virions of a same subtype is much more challenging due to their lower genetic diversity, and therefore remains understudied. We applied the methodology used by Kiwelu and colleagues [33] to the largest pure subtype datasets in ICONIC (subtypes B and C) and found similar intra-subtype recombination frequencies in subtypes B (13.4%) and C (9.1%).

Unfortunately, the low availability of HIV-1 full genomes in public databases prevented exploring more deeply the linkage of our dataset with non-UK epidemics, and also to provide more insights into the evolutionary history of the unique recombinants described here. Another limitation was the fact that not all our sequences were full-length genomes due to an unequal efficacy of the sequencing primers. In addition, although the study of HIV-1 strain distribution can provide valuable insights due to the different biological properties of certain subtypes, the growing heterogeneity of the viral epidemics will challenge the current HIV-1 strain classification.

In conclusion, the use of nearly full-genome sequences, as opposed to only PR+RT, provided a more reliable description of CRFs (that would be otherwise misclassified) and permitted the detection of novel recombinants circulating in London. Whereas it is likely that some of the URFs were imported from abroad, our work suggests that a large proportion of them could have been generated locally.

## Acknowledgements

We would like to thank the clinicians and scientists at University College London Hospital and Barts Health NHS Trust. We would also like to thank Dr Anna Tostevin (UCL Research Department of Infection and Population Health) for her help with the analysis of the UK HIV Drug Resistance Database sequences and Dr Astrid Gall (University of Cambridge) for her initial guidance on the HIV whole genome sequencing methods. This work was supported by the ICONIC project (supported by the Health Innovation Challenge Fund T5-344, a parallel funding partnership between the Department of Health and Wellcome Trust). The views expressed in this publication are those of the author(s) and not necessarily those of their funders and employees. The members of the ICONIC consortium are Dr A. Oliver, Dr C.Y. Tong, Dr D. Clark (Barts Health NHS Trust), Dr E. Nastouli (UCL Hospital NHS Foundation Trust), Prof A. Johnson, Prof D. Dunn, Prof A. Hayward, Prof D. Pillay, Prof S. Morris, Dr Z. Kozlakidis (UCL), Prof P. Kellam (Imperial College London), Prof J. Edgeworth (Guy’s and St. Thomas’ NHS foundation Trust) and Prof A. Leigh Brown (University of Edinburgh).

## References

1. LANL. Los Alamos HIV database 2017 [May 1, 2017]. Available from: http://www.hiv.lanl.gov.

2. LANL. HIV Circulating Recombinant Forms (CRFs) 2017 [May 1, 2017]. Available from: http://www.hiv.lanl.gov/content/sequence/HIV/CRFs/CRFs.html.

3. Hemelaar J, Gouws E, Ghys PD, Osmanov S. Global trends in molecular epidemiology of HIV-1 during 2000-2007. AIDS. 2011; 25: 679–89.

4. Beloukas A, Psarris A, Giannelou P, Kostaki E, Hatzakis A, Paraskevis D. Molecular epidemiology of HIV-1 infection in Europe: An overview. Infect Genet Evol. 2016.

5. Dolling D, Hué S, Delpech V, Fearnhill E, Leigh-Brown A, Geretti AM, et al. The increasing genetic diversity of HIV-1 in the UK, 2002-2010. AIDS. 2014; 28: 773–80.

6. Fernández-García A, Pérez-Álvarez L, Cuevas MT, Delgado E, Muñoz-Nieto M, Cilla G, et al. Identification of a new HIV type 1 circulating BF intersubtype recombinant form (CRF47_BF) in Spain. AIDS Res Hum Retroviruses. 2010; 26: 827–32.

7. Fernández-García A, Delgado E, Cuevas MT, Vega Y, Montero V, Sánchez M, et al. Identification of an HIV-1 BG Intersubtype Recombinant Form (CRF73_BG), Partially Related to CRF14_BG, Which Is Circulating in Portugal and Spain. PloS ONE. 2016; 11: e0148549.

8. Foster GM, Ambrose JC, Hue S, Delpech VC, Fearnhill E, Abecasis AB, et al. Novel HIV-1 recombinants spreading across multiple risk groups in the United Kingdom: the identification and phylogeography of Circulating Recombinant Form (CRF) 50_A1D. PloS ONE. 2014; 9: e83337.

9. Leoz M, Feyertag F, Charpentier C, Delaugerre C, Wirden M, Lemee V, et al. Characterization of CRF56_cpx, a new circulating B/CRF02/G recombinant form identified in MSM in France. AIDS. 2013; 27: 2309–12.

10. Simonetti FR, Lai A, Monno L, Binda F, Brindicci G, Punzi G, et al. Identification of a new HIV-1 BC circulating recombinant form (CRF60_BC) in Italian young men having sex with men. Infect Genet Evol. 2014; 23: 176–81.

11. Galimand J, Frange P, Rouzioux C, Deveau C, Avettand-Fenoel V, Ghosn J, et al. Short communication: evidence of HIV type 1 complex and second generation recombinant strains among patients infected in 1997-2007 in France: ANRS CO06 PRIMO Cohort. AIDS Res Hum Retroviruses. 2010; 26: 645–51.

12. Hemelaar J. Implications of HIV diversity for the HIV-1 pandemic. J Infect. 2013; 66: 391–400.

13. Mackie NE, Dunn DT, Dolling D, Garvey L, Harrison L, Fearnhill E, et al. The impact of HIV-1 reverse transcriptase polymorphisms on responses to first-line nonnucleoside reverse transcriptase inhibitor-based therapy in HIV-1-infected adults. AIDS. 2013; 27: 2245–53.

14. Peeters M, Aghokeng AF, Delaporte E. Genetic diversity among human immunodeficiency virus-1 non-B subtypes in viral load and drug resistance assays. Clin Microbiol Infect. 2010; 16: 1525–31.

15. Pant Pai N, Shivkumar S, Cajas JM. Does genetic diversity of HIV-1 non-B subtypes differentially impact disease progression in treatment-naive HIV-1-infected individuals? A systematic review of evidence: 1996-2010. J Acquir Immune Defic Syndr. 2012; 59: 382–8.

16. Kaleebu P, Nankya IL, Yirrell DL, Shafer LA, Kyosiimire-Lugemwa J, Lule DB, et al. Relation between chemokine receptor use, disease stage, and HIV-1 subtypes A and D: results from a rural Ugandan cohort. J Acquir Immune Defic Syndr. 2007; 45: 28–33.

17. Martinez-Cajas JL, Pant-Pai N, Klein MB, Wainberg MA. Role of genetic diversity amongst HIV-1 non-B subtypes in drug resistance: a systematic review of virologic and biochemical evidence. AIDS Rev. 2008; 10: 212–23.

18. deCamp A, Hraber P, Bailer RT, Seaman MS, Ochsenbauer C, Kappes J, et al. Global panel of HIV-1 Env reference strains for standardized assessments of vaccine-elicited neutralizing antibodies. J Virol. 2014; 88: 2489–507.

19. Zhuang J, Jetzt AE, Sun G, Yu H, Klarmann G, Ron Y, et al. Human immunodeficiency virus type 1 recombination: rate, fidelity, and putative hot spots. J Virol. 2002; 76: 11273–82.

20. Streeck H, Li B, Poon AF, Schneidewind A, Gladden AD, Power KA, et al. Immune-driven recombination and loss of control after HIV superinfection. J Exp Med. 2008; 205: 1789–96.

21. De Candia C, Espada C, Duette G, Ghiglione Y, Turk G, Salomon H, et al. Viral replication is enhanced by an HIV-1 intersubtype recombination-derived Vpu protein. Virol J. 2010; 7.

22. Ramirez BC, Simon-Loriere E, Galetto R, Negroni M. Implications of recombination for HIV diversity. Virus Res. 2008; 134: 64–73.

23. Moradigaravand D, Kouyos R, Hinkley T, Haddad M, Petropoulos CJ, Engelstadter J, et al. Recombination accelerates adaptation on a large-scale empirical fitness landscape in HIV-1. PloS genetics. 2014; 10: e1004439.

24. Moutouh L, Corbeil J, Richman DD. Recombination leads to the rapid emergence of HIV-1 dually resistant mutants under selective drug pressure. Proc Natl Acad Sci USA. 1996; 93: 6106–11.

25. Gall A, Ferns B, Morris C, Watson S, Cotten M, Robinson M, et al. Universal amplification, next-generation sequencing, and assembly of HIV-1 genomes. J Clin Microbiol. 2012; 50: 3838–44.

26. Robertson DL, Anderson JP, Bradac JA, Carr JK, Foley B, Funkhouser RK, et al. HIV-1 nomenclature proposal. Science. 2000; 288: 55–6.

27. Salminen MO, Carr JK, Burke DS, McCutchan FE. Identification of breakpoints in intergenotypic recombinants of HIV type 1 by bootscanning. AIDS Res Hum Retroviruses. 1995; 11: 1423–5.

28. Gallo Cassarino T, Frampton D, Sugar R, Charles E, Kozlakidis Z, Kellam P. High-throughput pipeline for de-novo assembly and drug resistance mutations identification from Next-Generation Sequencing viral data of residual diagnostic samples. 2016.

29. Struck D, Lawyer G, Ternes AM, Schmit JC, Bercoff DP. COMET: adaptive context-based modeling for ultrafast HIV-1 subtype identification. Nucleic Acids Res. 2015; 42: e144.

30. Schultz AK, Zhang M, Bulla I, Leitner T, Korber B, Morgenstern B, et al. jpHMM: improving the reliability of recombination prediction in HIV-1. Nucleic Acids Res. 2009; 37: W647–51.

31. Lole KS, Bollinger RC, Paranjape RS, Gadkari D, Kulkarni SS, Novak NG, et al. Full-length human immunodeficiency virus type 1 genomes from subtype C-infected seroconverters in India, with evidence of intersubtype recombination. J Virol. 1999; 73: 152–60.

32. Martin DP, Murrell B, Golden M, Khoosal A, Muhire B. RDP4: Detection and analysis of recombination patterns in virus genomes. Virus Evol. 2015; 1: vev003.

33. Kiwelu IE, Novitsky V, Margolin L, Baca J, Manongi R, Sam N, et al. Frequent intra-subtype recombination among HIV-1 circulating in Tanzania. PloS ONE. 2013; 8: e71131.

34. Hué S, Clewley JP, Cane PA, Pillay D. HIV-1 pol gene variation is sufficient for reconstruction of transmissions in the era of antiretroviral therapy. AIDS. 2004; 18: 719–28.

35. Brenner BG, Lowe M, Moisi D, Hardy I, Gagnon S, Charest H, et al. Subtype diversity associated with the development of HIV-1 resistance to integrase inhibitors. J Med Virol. 2011; 83: 751–9.

36. Zhao J, Tang S, Ragupathy V, Carr JK, Wolfe ND, Awazi B, et al. Identification and genetic characterization of a novel CRF22_01A1 recombinant form of HIV type 1 in Cameroon. AIDS Res Hum Retroviruses. 2010; 26: 1033–45.

37. Holguín A, Lospitao E, López M, de Arellano ER, Pena MJ, del Romero J, et al. Genetic characterization of complex inter-recombinant HIV-1 strains circulating in Spain and reliability of distinct rapid subtyping tools. J Med Virol. 2008; 80: 383–91.

